# Qualitative EEG abnormalities signal a shift towards inhibition-dominated brain networks. Results from the EU-AIMS LEAP studies

**DOI:** 10.1101/2024.09.19.613847

**Authors:** Arthur-Ervin Avramiea, Erika L. Juarez-Martinez, Pilar Garcés, Joerg F. Hipp, Simon-Shlomo Poil, Marina Diachenko, Huibert D. Mansvelder, Emily Jones, Luke Mason, Declan Murphy, Eva Loth, Bethany Oakley, Tony Charman, Tobias Banaschewski, Bob Oranje, Jan Buitelaar, Hilgo Bruining, Klaus Linkenkaer-Hansen

**Affiliations:** Department of Integrative Neurophysiology, Center for Neurogenomics and Cognitive Research, (CNCR), Amsterdam Neuroscience, VU Amsterdam, 1081 HV Amsterdam, The Netherlands; Child and Adolescent Psychiatry and Psychosocial Care, Emma Children’s Hospital, Amsterdam UMC, Vrije Universiteit Amsterdam, Meibergdreef 5, 1105 AZ, Amsterdam, The Netherlands; Roche Pharma Research and Early Development, Neuroscience and Rare Diseases, Roche Innovation Center Basel, Basel, Switzerland; Aspect Neuroprofiles BV, Amsterdam, The Netherlands; Department of Psychological Sciences, Centre for Brain and Cognitive Development, Birkbeck, University of London, London, UK; Department of Forensic and Neurodevelopmental Sciences, Institute of Psychiatry, Psychology and Neuroscience, King’s College London, UK; Institute for Translational Neurodevelopment/ South London and Maudsley NHS Foundation Trust; Department of Psychology, Institute of Psychiatry, Psychology and Neuroscience, King’s College London, UK; Department of Child and Adolescent Psychiatry and Psychotherapy, Central Institute of Mental Health, Medical Faculty Mannheim, University of Heidelberg, German Center for Mental Health (DZPG), partner site Mannheim-Heidelberg-Ulm, Mannheim, Germany; Center for Neuropsychiatric Schizophrenia Research (CNSR), Copenhagen University Hospital, Psychiatric Center Glostrup, Denmark; Department of Cognitive Neuroscience, Donders Institute for Brain, Cognition and Behaviour, Radboudumc, Nijmegen, The Netherlands; N=You Neurodevelopmental Precision Center, Amsterdam Neuroscience, Amsterdam Reproduction and Development, Amsterdam UMC, Amsterdam, The Netherlands; Levvel, Center for Child and Adolescent Psychiatry, Amsterdam, The Netherlands

## Abstract

Qualitative EEG abnormalities are common in Autism Spectrum Disorder (ASD) and hypothesized to reflect disrupted excitation/inhibition balance. To test this, we recently introduced a functional measure of network-level E/I ratio (fE/I). Here, we applied fE/I and other EEG measures to alpha oscillations from source-reconstructed data in the EU-AIMS dataset (267 ASD, 209 controls). We analyzed these measures alongside qualitative EEG abnormalities ranging from slowing of activity to epileptiform patterns, aiming to replicate the findings from the SPACE-BAMBI study. EEG abnormalities were rare in adults and could not be statistically assessed. ASD children-adolescents with EEG abnormalities exhibited lower relative alpha power and fE/I compared to those without. However, EEG-abnormality scoring did not stratify the behavioral heterogeneity of ASD using clinical measures. Surprisingly, several controls also exhibited qualitative EEG abnormalities with a strikingly similar anatomical distribution of reduced fE/I, suggesting a shift towards inhibition-dominated network dynamics in sensory processing regions. The robustness of this association between EEG abnormalities and reduced fE/I was further supported by re-analysis of the SPACE-BAMBI study in source space. Stratification by the presence of EEG abnormalities and their effects on network activity may help understand neurodevelopmental physiological heterogeneity and the difficulties in implementing E/I targeting treatments in unselected cohorts.

## Introduction

The opposing forces of excitation (E) and inhibition (I) fundamentally shape activity at many levels of neuronal organization. Heuristic arguments have favored a certain balance in E/I ratios to be important for normal brain function^1^, which has been underscored by a large body of computational modeling in recent years^2–5^. Disturbances in E/I ratios at various levels of neuroscientific interrogation have been associated with autism spectrum disorder (ASD)^6–17^, among other disorders^18^. The original implication of E/I imbalances in ASD was based upon its high comorbidity with epilepsy and the frequently observed EEG abnormalities (ranging from epileptiform patterns to non-epileptiform activity such as slowing)^19–22^. Preclinical support for the E/I imbalance theory ranges from genetic studies on synaptic proteins to postmortem analyses of GABAergic and glutamatergic systems^11,12,23–25^. Collectively, the results indicate that up and down regulation of both excitation and inhibition may contribute to ASD manifestations, probably reflecting a mixture of primary and secondary interactions around homeostatic mechanisms^25^. Thus, variation in E/I directionality within ASD may result in different forms of pathophysiology and, in turn, variability in treatment response^17,24,25^.

An important challenge is to develop clinically-applicable methods to increase our understanding of global network E/I imbalances in addition to the knowledge gained at the local or cellular level. In vivo, markers of E/I balance are being developed, e.g., using magnetic resonance spectroscopy, which can quantify concentrations of inhibitory (GABA) and excitatory (glutamate) neurotransmitters^13–17^. Although promising, these studies are hampered by the limited spatial and temporal resolution. More recently, quantitative EEG techniques have been proposed as proxy markers for E/I balance in mass brain activity^26^. Our group has put forward a definition of network E/I balance that is inspired by computational modeling of critical brain dynamics^27,28^. In this framework, E/I balance is an emergent network property—characterized by high spatio-temporal complexity emerging in neuronal networks balancing between order and disorder. This definition enables measuring a network-level functional E/I ratio (fE/I), which is useful for understanding basic principles of information processing in neuronal networks^2–4,29–31^ and for studying brain disorders in which E/I balance may be disrupted. The fE/I method has shown sensitivity to changes in both network connectivity and synapse functioning through simulations^3^. Importantly, it is directly applicable to EEG recordings, making it well-suited for clinical use. Indeed, fE/I estimation in EEG recordings corresponding to a large sample of healthy controls, confirmed an overall balanced network E/I ratio. We further validated the method using pharmacological manipulation in healthy subjects with a GABAergic drug^27^.

In parallel, our group investigated network-level fE/I for the first time in a clinical population of children with ASD^27^. In line with the aforementioned findings from preclinical studies, we found that ASD is characterized by both increased and decreased E/I ratios supporting the notion that E/I variability is substantial. Importantly, we found a strong association between qualitative EEG abnormalities identified through visual inspection and fE/I variability, supporting the hypothesis that epileptiform activity and EEG abnormalities relate to E/I imbalances. Furthermore, we implemented fE/I in an innovative randomized controlled trial testing bumetanide as an E/I targeting treatment and confirmed that the measure is treatment sensitive^32^. These findings provide an initial proof-of-concept of the utility of fE/I estimation for physiological stratification in ASD^27^.

Based on these initial observations, we hypothesize that establishing an association between qualitative and quantitative EEG abnormalities in ASD is instrumental to understand the implication of E/I imbalances towards network and information processing dysfunctions. In this line, fE/I estimation comes to light as an appropriate measure to study the diverse network-level E/I dynamics that may result from primary pathophysiological mechanisms or compensatory mechanisms that counteract epileptogenic activity. Here, we elaborated on these hypotheses and investigated fE/I variability in the large EU-AIMS compilation of EEG recordings in children-adolescents and adult samples with or without ASD. We make three predictions: first, that the EU-AIMS dataset reproduces the increased variability in fE/I observed in ASD when compared to controls^27^; second, that qualitative EEG abnormalities are associated with alterations in network E/I as reported in^27^; and third, that the presence of qualitative EEG abnormalities modulates the effect of network E/I on clinical outcome.

## Materials and methods

### Participants

#### EU-AIMS

Participants aged 6 to 32 years were recruited between January 2014 and March 2017 across five European specialist ASD centers and their EEG data collected as part of the EU-AIMS Longitudinal European Autism Project (LEAP). A complete description of the study design and clinical characterization of the participants can be found in^33,34^. Inclusion criteria for the ASD sample were an existing clinical diagnosis of ASD according to the DSM-IV^35^, DSM-IV-TR, DSM-5^36^, or ICD-10^37^ criteria. ASD symptoms were additionally assessed using the Autism Diagnostic Observation Schedule (ADOS^38,39^) and the Autism Diagnostic Interview-Revised (ADI-R^40^). However, individuals with a clinical ASD diagnosis who did not reach cut-offs on these instruments were not excluded. Psychiatric conditions (except for psychosis or bipolar disorder) and medication use were allowed. Exclusion criteria for the control sample were a T score of ≥ 70 on the self-report or parent-report form of the Social Responsiveness Scale^41^, presence of any psychiatric disorder and/or psychoactive medication use, and a total IQ <75 (Intellectual disability). For the ASD sample in this study, an IQ < 50 was an exclusion criteria to include a wide range of the ASD spectrum but also limit it to the criteria used in the earlier study that we aim to replicate^27^. The clinical scales included were the Repetitive Behavior Scale-Revised (RBS-R; total raw score; range 0−129, higher score indicates more affected)^42^ and the Vineland-II Adaptive Behavior standard score (VABS^43^). The latter assess the adaptive behavior grouped into four major domains: Communication, Daily living, Socialization and Motor skills (VABS-com, VABS-dl and VABS-soc respectively). Sums of the scores from these domains are standardized, higher scores indicate better adaptive behavior (range 86–100) and lower scores indicate low adaptive behavior (<85). Finally, the sums of the domain standard scores are further standardized into an Adaptive Behavior Composite Score (VABS-composite). The study was approved by the medical ethical committee at each site independently. All participants or their legal guardians signed informed consent.

#### SPACE-BAMBI

EEGs and symptom-scale baseline measurements were collected from two ongoing studies at the developmental disorder unit in the UMC Utrecht, with identical EEG measurement protocols (SPACE (Sensory information Processing in Autism and Childhood Epilepsy) and BAMBI (Bumetanide in Autism Medication and Biomarker, Eudra-CT 2014-001560-35)). Inclusion criteria for ASD and TDC samples were an age between 7–16 years, IQ > 55, and ability to comply with study procedures. ASD diagnosis required an expert diagnosis according to the Diagnostic and structural manual of mental disorders (DSM) IV-TR^35^ or 5^36^ supported by a clinical score on the Autism Diagnostic Observation Schedule 2 (ADOS-2 module 3 or 4, score ≥ 7) or a subclinical score on the Social Responsivity Scale (SRS-t-score ≥ 60)^44^. An abbreviated form of the Wechsler intelligence scale for children (WISC)-III was used for IQ estimation. Exclusion criteria for the ASD sample were use of psychoactive medication and presence of other major neurological disease such as previous or current epilepsy diagnoses. The TDC sample was recruited from local residents attending non-special education, where a history of behavioral or learning problems, a diagnosis of any neurodevelopmental condition or any other major health issue was an exclusion criterion.

We recruited a total of 126 subjects that met the inclusion criteria for ASD and 37 TDC. From these, 26 children-adolescents with ASD and 8 TDC were excluded because less than 50% of their recordings were free of transient artifacts or no eyes-closed rest EEG had been recorded. Thus, the final included sample consisted of 129 subjects (42 females): 100 with ASD (7–16 years, M = 10.5 years, SD = 2.3 years, 27 females) and 29 age-matched typically developing children-adolescents (TDC) (7.4–14.4 years, M = 10.3, SD = 1.5 years, 15 females). The study was approved by the medical ethical committee. All participants or their legal guardians signed informed consent.

### EEG recordings and pre-processing

#### EU-AIMS

EEGs were recorded during 4 minutes of resting state per participant (2 min eyes open, 2 min eyes closed) in five different centers: Central Institute of Mental Health (CIMH, Mannheim, Germany), Institute of Psychiatry, Psychology and Neuroscience, King’s College London (KCL, UK), Radboud University Nijmegen Medical Centre (RUNMC, Netherlands), University Campus BioMedico (UCBM, Rome, Italy) and University Medical Centre Utrecht (UMCU, Netherlands). The following EEG systems were employed: Brain Vision (CIMH, KCL, RUNMC), BioSemi (UMCU) and Micromed (UCBM), with sampling frequencies of 5000 Hz (KCL, RUNMC), 2048 Hz (UMCU), 2000 Hz (CIMH) and 256–1000 Hz (UCBM). All sites used 10-20 layout caps, with 60–64 electrodes. To optimize participant compliance, resting state was acquired in 30 sec blocks, alternating eyes open (fixating an hourglass) and eyes closed. Segments were then concatenated into a single signal for each condition. To simplify our analysis and in accordance with Bruining *et al.*^27^, only the eyes-closed-rest (ECR) condition was analyzed for this study, since, in children, ECR is less prone to movement and other artifacts. Researchers were blind to group label and the clinical characteristics of the participants (other than age) until after the EEG analysis was carried out (Neurophysiological Biomarker Toolbox and custom-made scripts^45^). Data were homogenized by selecting 61 common electrodes (data from outer electrodes FT9, FT10, TP9, TP10, P9, P10, PO9, PO10 and Iz, if recorded, was discarded, since these electrodes were often noisy or not standard in all the samples), resampled to 1000 Hz and band-pass filtered to 1–32 Hz with a finite impulse response filter. Artifact rejection was performed manually: we eliminated the bad channels and then discarded noisy intervals, after which we re-referenced to the average reference. EEGs were considered of sufficiently good quality to reliably estimate temporal and excitation-inhibition ratio biomarkers (*see EEG Analysis below*), if they contained enough noise-free segments of ≥2 seconds to add up to at least 80 seconds of data. Following this criteria, 63 recordings were excluded from analysis (control group: *n* = 19, ASD group: *n* = 44; **Figure 1**). After pre-processing, on average 108 seconds per recording (84–131 s) were available for analysis for the ASD group. For the control group on average 110.3 seconds per recording (80–132 s) were available for analysis.

**Figure 1.**
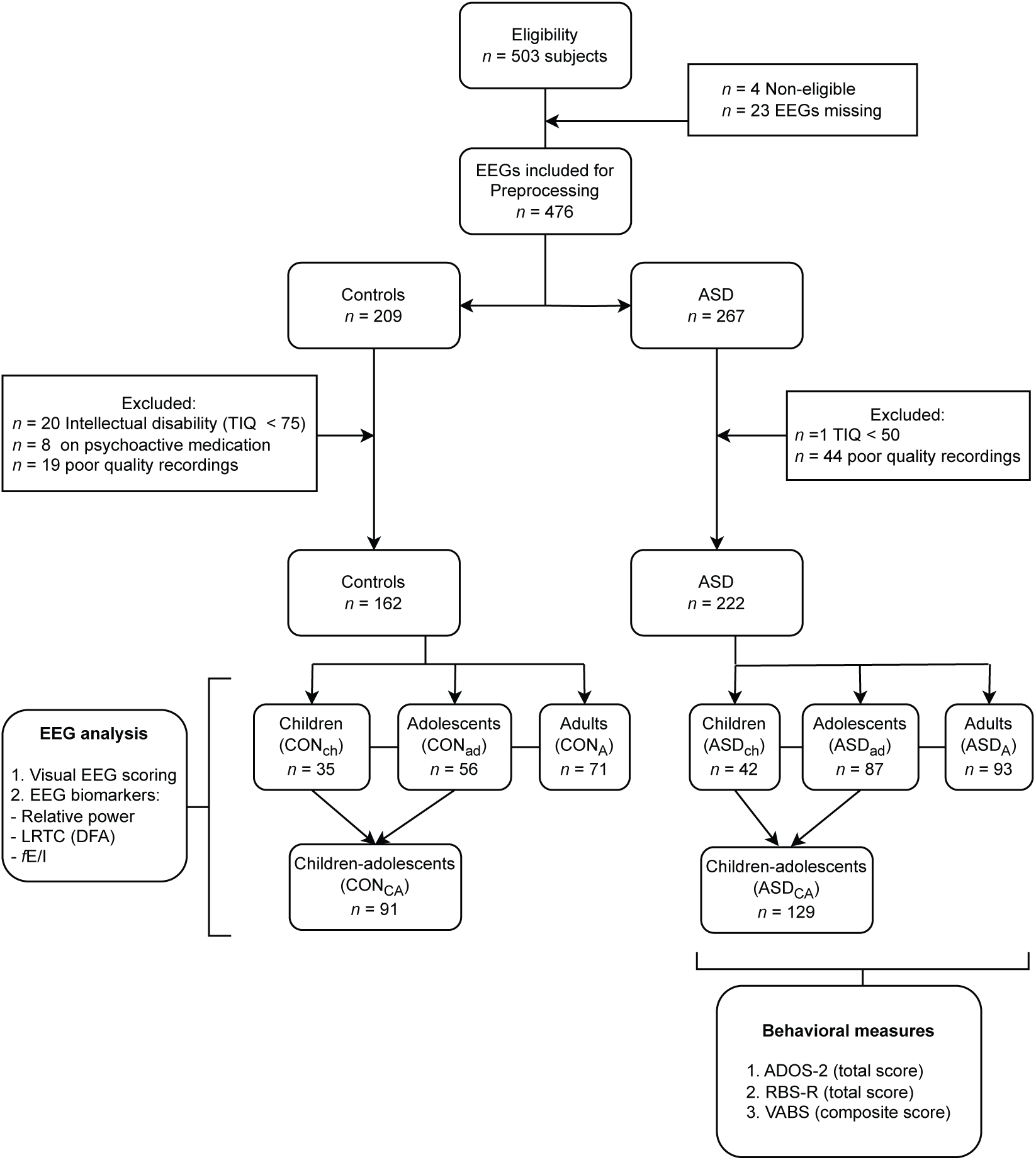
Flow diagram of the participants included for EEG analysis from the EU-AIMS Longitudinal European Autism Project (LEAP). A total of 503 subjects were screened for eligibility between January 2014 and March 2017 across five European specialist ASD centers. EEGs from 476 subjects were available. After preprocessing and exclusion criteria, 162 EEGs were included in the control group and 222 EEGs in the ASD group. All EEGs were cleaned, visually scored, and the following biomarkers were computed in the alpha-frequency band: relative power, Long-range temporal correlations (LRTC) quantified by detrended fluctuation analysis (DFA), and functional excitation-inhibition ratio (fE/I). Behavioral measures were available for the ASD group: Repetitive Behavior Scale-Revised (RBS-R); Vineland-II Adaptive Behavior standard scores (VABS); VABS Composite standard score (VABS-composite).

#### SPACE-BAMBI

EEGs were recorded during 3–5 minutes of eyes-closed rest at the UMC Utrecht using A 64-channel BioSemi EEG system at a sampling rate of 2048 Hz and common mode sense (CMS) reference electrode.

EEG analyses were initially pre-processed using the in-house developed Neurophysiological Biomarker Toolbox (NBT) written in MATLAB. All recordings were manually cleaned for artifacts, i.e., noisy channels were discarded and noisy intervals removed. Subsequently, the data were re-referenced to the average reference. An average of 217 seconds (range 61–308 s) per recording were available for analysis for the children with ASD and TDC samples.

### EEG source localization

For source reconstruction of the EEG signals, we used the pipeline described in Li *et al.*^46^. To obtain cortical current estimates from our sensor level data, we used the MNE-Python implementation of the L2 minimum norm estimation. To construct the boundary element head model and forward operator for the source modelling, we used the FreeSurfer average brain template from FreeSurfer 6^47^. The inverse operator was constructed using the default MNE parameters, and was applied with the regularization parameter set to *λ*^2^ = 1/9. Unconstrained orientations were allowed, and principal component analysis was applied to the source time series at each vertex to reduce the three-dimensional signals to one-dimensional time series of the dominant principal component. The time series from the 20484 source vertices were further collapsed into 68 cortical patches using the Desikan Killiany atlas. This process involved aligning the dipole orientations by shifting vertices with opposite polarity to the majority of vertices by *π*, and then averaging the amplitudes of all vertices within a patch. The phase shifting ensured that vertices with opposite polarities did not cancel each other out during the averaging operation.

### EEG analysis

We focused our analysis on alpha-band oscillations (8–13 Hz) due to their relevance for healthy neuronal network development and cognitive function^48^, their clear disruption in neurodevelopmental disorders, but also to reduce the number of comparisons^49,50^.

#### Power

Spectral power was computed using multitaper spectral estimation^51^. Relative alpha power is expressed in percent and was calculated by dividing the absolute power in the alpha band with the integrated power in the range of 1–45 Hz.

#### Temporal structure

Neuronal oscillations exhibit complex fluctuations in amplitude that are characterized by a power-law decay of auto-correlations, also known as LRTC^52^. Computational modeling has shown that the strength of LRTC is influenced by the balance between excitatory and inhibitory signaling in the network producing the oscillations^28^. We used the detrended fluctuation analysis (DFA) to quantify LRTC in the amplitude modulation of alpha oscillations in the time range of seconds to tens of seconds^53^. The amplitude envelope of the alpha oscillations was extracted using the Hilbert transform applied to the FIR-filtered signals in the 8–13 Hz band. The analytical steps to quantify LRTC using DFA have been explained in detail previously^45,52^. Here, we used a standard assessment of the strength of LRTC on time scales from 2 to 20 seconds, which is reflected in the so-called “DFA exponent”. An exponent of 0.5 characterizes an uncorrelated signal whereas an exponent in the interval of 0.5 to 1.0 indicate long-range temporal correlations with larger exponents indicating stronger correlations. in the alpha frequency band (8–13 Hz).

#### Excitation/inhibition ratio (fE/I)

The algorithm for estimating E/I ratio, was derived from the Critical Oscillations (CROS) model of ongoing neuronal activity^28^, where a strong association is observed between the ratio of excitation/inhibition connectivity, oscillation amplitude, and LRTC. From these observations, a functional form of E/I ratio (*fE*/*I*) is estimated based on the Pearson correlation between the amplitude and LRTC within a signal. This method was validated using pharmacological manipulation in healthy subjects using a GABAergic drug (zolpidem), corroborating a reduction of *f*E/I ratios after the drug administration, and already tested in an ASD sample supporting the notion of large heterogeneity in E/I ratios (for details see Bruining *et al.*^27^). The analytical steps for the algorithm are as follows: to test the relationship between the amplitude and LRTC of the amplitude envelope of an oscillatory signal, it is necessary to have a measure of LRTC on short time-scales that is unbiased by the amplitude of the signal. To this end, an amplitude-normalized fluctuation function, *nF*(*t*), is calculated as follows: The signal is band-pass filtered (*i*), the amplitude envelope *A* extracted (*ii*), the signal profile, *S* can then be calculated as the cumulative sum of the demeaned amplitude envelope (*iii*),

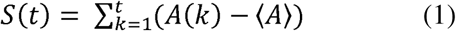

and split into windows of a certain size (the default used here is 5 seconds) in exactly the same way as during the DFA calculation^45,53^. As an additional step (*iv*), each of these signal-profile windows is divided by the mean of the amplitude envelope for that window calculated during step (*ii*). These amplitude-normalized windows are then detrended (*v*) and, subsequently, the normalized fluctuation function is calculated for each window as the standard deviation of the amplitude-normalized signal profile (*vi*). To calculate the functional excitation/inhibition ratio, *fE*/*I*, Pearson correlation between the amplitude and the normalized fluctuation function for the set of windows *W* (*vii*) is performed. *fE*/*I* is then defined as:

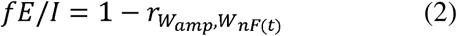

Sub-critical (inhibition-dominated) networks will have an *fE*/*I* < 1, super-critical (excitation-dominated) networks *fE*/*I* > 1, and critical (E/I-balanced) networks will have *fE*/*I* = 1. A DFA > 0.6 inclusion criterion for networks or channels before computing the *fE*/*I* is used because networks without LRTC will not show a co-variation of amplitude and the fluctuation function. For our analyses, *fE*/*I* was calculated for windows of 5 seconds with 80% overlap.

### Classification of EEG abnormalities

All recordings were visually inspected in windows of 10 seconds by a neurologist with training in clinical EEG (neurophysiology and epilepsy; EJM) and whom remained blind to the group label, and scored according to Luders & Noachtar’s classification of EEG abnormalities^54^. When no abnormalities were detected, the EEG was classified as normal.

### Statistical analysis

Two age categories were defined: children-adolescents (age 6–17 years) and adults (age 18– 31 years). For both age groups, whole-brain average EEG biomarker values, as well as biomarker values over the 68 source-localized patches, were compared between the control groups (i.e., CON_C_: control children-adolescents; CON_A_: control adults) and the ASD groups (i.e., ASD_C_: children-adolescents with ASD; ASD_A_: Adults with ASD). Finally, we investigated the correlation between EEG measures and clinical scores for the ASD groups.

### Statistical analysis within EEG-abnormality subgroups

Participants with ASD were grouped based on the presence or absence of EEG abnormalities (ASD_C-qNL_, ASD_C-qABN_; ASD_A-qNL_, ASD_A-qABN_) and compared against the control groups and against each other (whole-brain average and per region). Additionally, we investigated the correlation between EEG measures in each subgroup and the clinical scores.

Non-specific EEG abnormalities can be present in the normal population. Although their clinical meaning is debated, they could be markers of common neurological conditions or unrecognized psychiatric disorders^55–58^. In this study we found EEG abnormalities in 9.8% of the subjects included in the control groups (CON_C-qABN_, CON_A-qABN_). To investigate if they showed distinct clinical and EEG characteristics, we compared them to controls with normal EEG (CON_C-qNL_, CON_A-qNL_), and also to the ASD_qABN_ subgroups. We also determined the difference between controls and ASD when no EEG abnormalities were present.

All patch-level comparisons were performed using Wilcoxon rank-sum test. For whole-brain analyses, we controlled for age, sex and IQ as covariates (ANCOVA test) and reported the adjusted *p-*values. Correlations between EEG measures and clinical scores were performed using partial (Pearson or Spearman) correlation coefficient, depending on whether normality assumptions were met, to additionally control for age, sex and IQ^59^. For patch-level analyses, False Discovery Rate (FDR, *q* = 0.05) was used to correct for multiple testing. Significance level was set at *p* < 0.05.

## Results

503 participants aged 6–32 years old were recruited across five European specialist ASD centers and their EEG data collected as part of the EU-AIMS Longitudinal European Autism Project (LEAP)^33,34^. EEG recordings of 476 subjects were available. Given differences in EEG development throughout the lifespan^60–65^, the data were divided into three age groups: children (6–11 years), adolescents (12–17 years), and adults (18–31 years). Due to the low sample sizes for the children and adolescents groups (**Figure 1**), and in order to simplify our analysis, we merged these into a single children-adolescents group (6–17 years) similar to Bruining *et al.*^27^. After exclusion criteria and EEG preprocessing (**Figure 1**), the control groups comprised 91 children-adolescents (CON_CA_, 7–17 years, *M* = 13.0 ± 3.0 years, 34 females) and 71 adults (CON_A_, 18–31 years, *M* = 22.6 ± 3.6 years, 19 females). The ASD groups comprised 129 children-adolescents (ASD_CA_, 6–17 years, *M* = 13.0 ± 3.0, 38 females) and 93 adults (ASD_A,_ 18–30 years, *M* = 21.8 ± 3.2, 21 females). Demographic and main clinical characteristics of the sample included are described in **Table S1.** There was no difference in age between groups. As expected, the mean total IQ for the CON_C_ and CON_A_ was higher than the ASD_CA_ and ASD_A_, respectively (CON_CA_: *M_IQ_* = 109.8 ± 1.4, ASD_CA_: *M_IQ_* = 98.3 ± 1.7, *p* <.0001; CON_A_: *M_IQ_* = 108 ± 1.4, ASD_A_: *M_IQ_* = 99.6 ± 1.9, *p* = .005) (**Table S1**).

In these EEG samples, we investigated the relative spectral power, long-range temporal correlations (LRTC, quantified by the DFA exponent), and functional excitation/inhibition ratio (fE/I) of alpha oscillations recorded with EEG during 2 minutes of eyes-closed rest.

### ASD and controls exhibit similar variability of network-level E/I

First, we tested whether the current dataset replicates the previous finding of higher variability in ASD subjects compared to controls^27^. We did not observe significant EEG differences in mean or variance of alpha-band relative power, DFA exponent, or fE/I in either the children with ASD (ASD_ch_) or adolescent groups (ASD_ad_), at regional or at whole-brain level when compared to controls (**Figure S1, Table S2**). The lack of differential effects between children and adolescents in terms of alpha-band relative power, LRTC and fE/I, supported us merging these groups into one age group for greater statistical power in subsequent analyses (**Figure S2**). This age merge (children-adolescents, CA) did not change the results, and EEG differences remained non-significant for all compared biomarkers (**Figure 2**), both at patch and at whole-brain level (*relative power* (mean ± SEM): CON_CA_= 0.33 ± 0.01, ASD_CA_= 0.30 ± 0.01, *p* = 0.34, *p_Levene_* = 0.28; *DFA*: CON_CA_= 0.68 ± 0.01, ASD_CA_= 0.67 ± 0.01, *p* = 0.52, *p_Levene_* = 0.14; *fE/I*: CON_CA_= 1.01 ± 0.02, ASD_CA_= 0.99 ± 0.01, *p* = 0.26, *p_Levene_* = 0.24).

**Figure 2.**
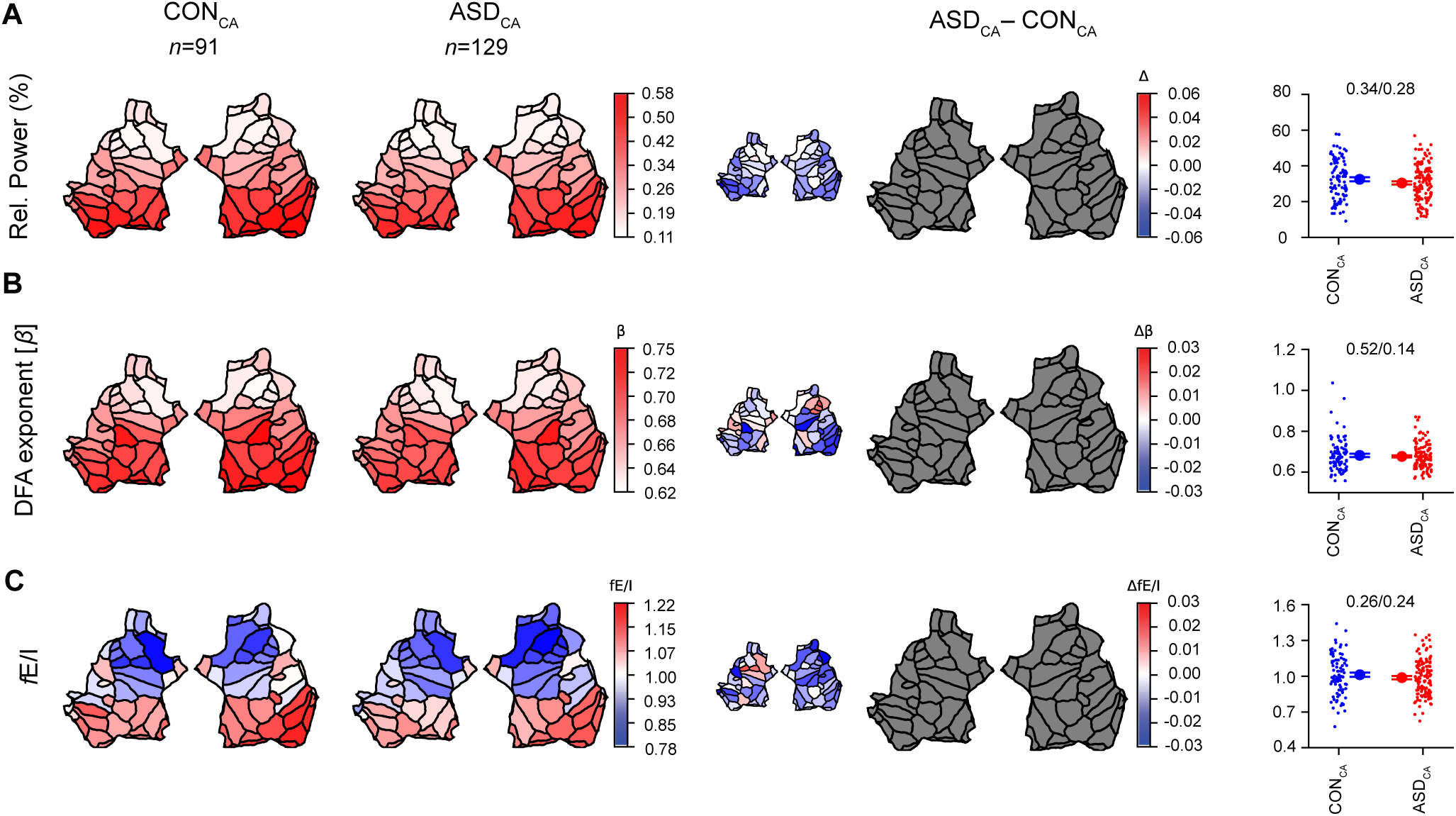
Alpha oscillations are quantitatively similar in the unstratified children-adolescents groups of ASD and controls. Grand-average flatmap representations of the two hemispheres, consisting of 68 patches (sources in the brain), are shown for CON_CA_ (*first column*) and ASD_CA_ (*second column*) for the alpha-band biomarkers. The grand-average difference plot of ASD_CA_-CON_CA_ is shown in the small (*3^rd^ column*), whereas the large plot represents significant differences after FDR correction as colored patches, and insignificant differences as gray patches (*4^th^ column*). There were no significant differences in mean or variance of **(A)** relative power, **(B)** DFA, or **(C)** fE/I in the alpha band, between groups at the patch level (*4^th^ column*) nor at the whole-brain level (*5^th^ column*). Comparisons in boxplots (*5^th^ column*) were based on the average value of the EEG biomarkers across all 68 patches (*p*-values are from ANCOVA / Levene’s test of variability). Individual-subject values, mean, and SEM are shown for CON_CA_ (*blue circles*) and ASD_CA_ (*red circles*). For all comparisons at the whole-brain level, *p*-values are adjusted for age, IQ, and sex as covariates.

Adults with ASD, however, showed wide-spread lower alpha-band relative power, and lower fE/I in frontal regions compared to controls (**Figure S2, Table S2)**. The lower fE/I in ASD adults was also significant at the whole-brain level (**Figure S2, Table S2**). Of note, neither of the age groups with ASD presented with larger fE/I variance compared to controls in this sample.

### Qualitative EEG abnormalities are associated with quantitative EEG differences in children-adolescents with ASD and controls

Following our previous observation that E/I variability may be related to qualitative EEG abnormalities^27^, we graded the EEGs for qualitative abnormalities using the EEG classification of Lüders & Noachtar^54^ (Methods), and investigated their association with spectral power, DFA exponents, and fE/I whilst being blinded to the group label and clinical characteristics of the participants.

In the children-adolescents group, we identified qualitative EEG abnormalities in 17% of the ASD_CA_ cases (ASD_CA-qABN,_ *n* = 22), ranging from slowing of activity to epileptiform abnormalities. Of note, none of the subjects had a clinical history of seizures or had been diagnosed with epilepsy. We also found EEG abnormalities in 14% of the children-adolescents included in the CON_CA_ group (*n* = 13). In terms of clinical and demographical characteristics, children-adolescents with qualitative EEG abnormalities in both subgroups were significantly younger than those without abnormalities (CON_CA-qNL_, *M* = 13.4 ± 2.9 years vs. CON_CA-qABN_, *M* = 10.8 ± 2.5 years, *p* = .006; ASD_CA-qNL_, *M* = 13.3 ± 2.9 years vs. ASD_CA-qABN_, *M* = 11.5 ± 3.0 years; *p* = .006) (**Table S3**). While the presence or absence of qualitative EEG abnormalities did not have an impact on the clinical scales in children-adolescents with ASD, typically developing children-adolescents with qualitative EEG abnormalities showed a significantly higher VABS-composite and VABS-communication scores (**Table S3**).

In the adults group, we identified qualitative abnormalities in only 5% of the ASD_A_ cases (ASD_A-qABN_, *n* = 5), and in 4% of the CON_A_ subjects (CON_A-qABN_, *n* = 3). Due to the low numbers in the adults group, we could not investigate quantitative EEG differences between subgroups defined by qualitative EEG abnormalities.

Children-adolescents with ASD and abnormal EEG (ASD_CA-qABN_, *n* = 22) had significantly lower alpha-band relative power, when compared to children-adolescents with normal EEG (ASD_CA-qNL_, *n* = 107), in all but one patch (**Figure 3A**), and also at the whole-brain level (mean ± SEM: ASD_CA-qNL_= 0.33 ± 0.01, ASD_CA-qABN_= 0.18 ± 0.01, *p* < .0001, *p_Levene_* = 0.028). fE/I was significantly lower in children with abnormal EEG, with the effects concentrated in the occipital region, but also present in the temporal, cingulate and frontal regions pertaining to the right hemisphere (**Figure 3C, Table S4**). A significant reduction in fE/I with abnormal EEG was also observed at the whole-brain level (mean ± SEM: ASD_CA-qNL_ = 1.003 ± 0.014, ASD_CA-qABN_ = 0.91 ± 0.031, *p* = 0.01, *p_Levene_* = 0.66). No significant changes were observed in DFA (mean ± SEM: ASD_CA-qNL_ = 0.67 ± 0.01, ASD_CA-qABN_ = 0.67 ± 0.01, *p* = 0.90, *p_Levene_* = 0.69). We also found significantly lower alpha-band relative power in the control children-adolescents with abnormal EEG (CON_CA-qABN_, *n* = 13) compared to CON_CA-qNL_ (*n* = 78, **Figure 3D**), both at patch and at whole brain-level (mean ± SEM: CON_CA-qNL_= 0.35 ± 0.01, CON_CA-qABN_= 0.17 ± 0.01, *p* < .0001, *p_Levene_* = 0.005). fE/I in control-adolescents with abnormal EEG was lower at the whole-brain level (mean ± SEM: CON_CA-qNL_ = 1.04 ± 0.02, CON_CA-qABN_ = 0.86 ± 0.04, *p* = 0.0007, *p_Levene_* = 0.55), was present in patches localized in the bilateral occipital regions, but also bilateral temporal and cingulate regions, right parietal and insular cortex as well as the left frontal region (**Figure 3F, Table S5**). No significant changes were observed in DFA (mean ± SEM: CON_CA-qNL_ = 0.68 ± 0.01, CON_CA-qABN_ = 0.68 ± 0.02, *p* = 0.60, *p_Levene_* = 0.67). Notably, subjects with qualitative EEG abnormalities in the control (CON_CA-qABN_) and ASD groups (ASD_CA-qABN_) had comparable alpha-band relative power, DFA exponent, and fE/I (**Figure 4A-C**), as did subjects in the ASD_CA-qNL_ and CON_CA-qNL_ groups (**Figure 4D-F, Table S6**). No difference in DFA was observed between any of the groups **(Figure 3B, E, Figure 4B, E).**

**Figure 3.**
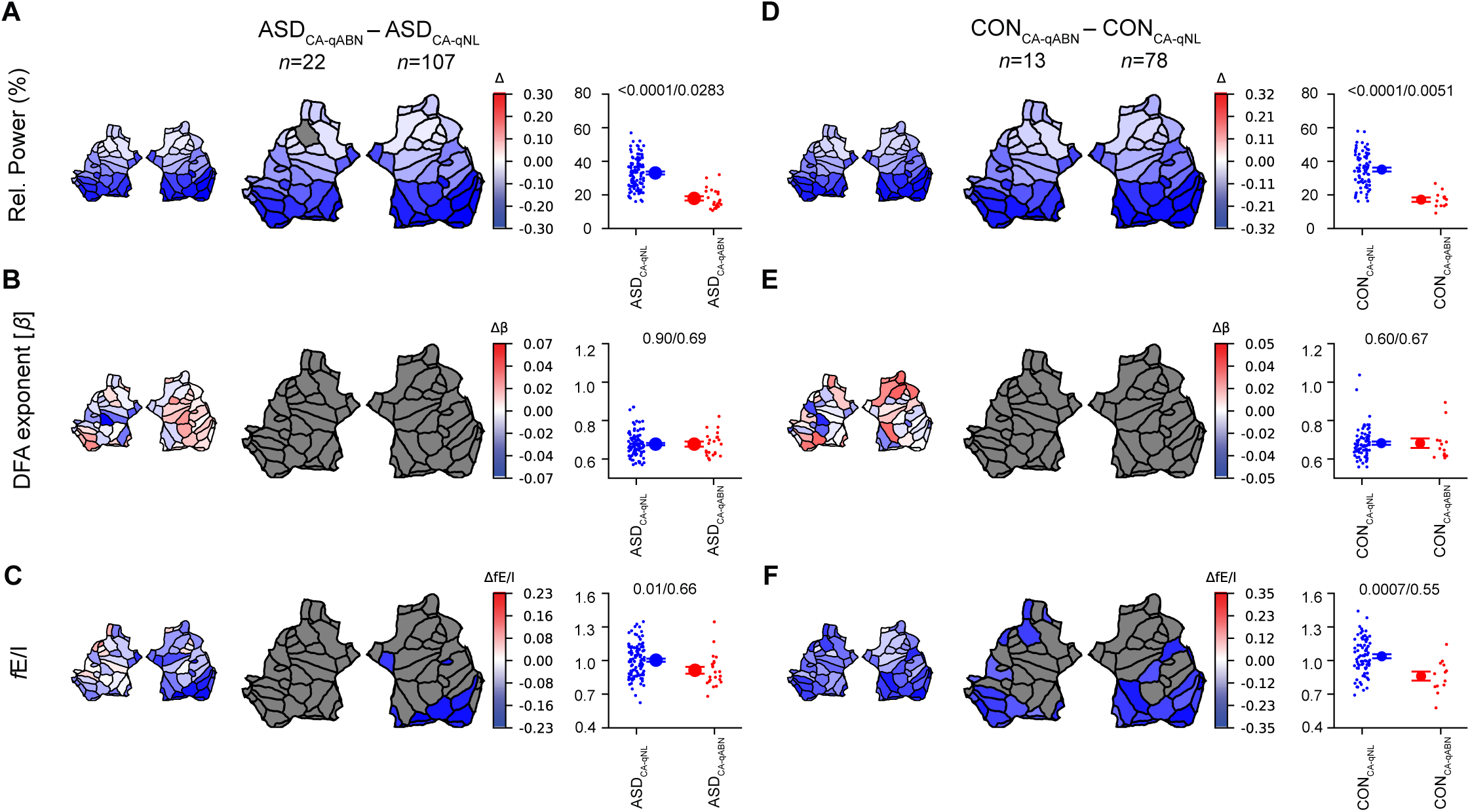
Children-adolescents with qualitative EEG abnormalities in both ASD and control groups show lower relative power and lower fE/I in the alpha-band, compared to those with normal EEG. Children-adolescents with ASD and abnormal EEG (ASD_CA-qABN_) had lower alpha-band relative power **(A)** and lower fE/I mean **(C)** compared to ASD children-adolescents with normal EEG (ASD_CA-qNL_). Interestingly, control children-adolescents with abnormal EEG (CON_CA-qABN_) also showed lower relative alpha power **(D)** and lower fE/I **(F)** compared to controls with normal EEG (CON_CA-qNL_) in similar brain regions as the ASD group. No difference in DFA was observed between groups **(B, E)**. Relative alpha power exhibited a significantly narrower range of values both in the abnormal group of ASD and controls compared to the groups with qualitatively normal EEG (Levene’s test) **(A, D)**. Grand-average flatmaps are shown for the cohort difference of the indicated comparisons (labels on top), where the small flatmaps represents the mean difference, and the large plot represents significant differences (*p*-value < 0.05, Wilcoxon rank-sum test, FDR corrected) as colored patches, and insignificant differences as gray patches. Comparisons in boxplots were based on the average value of the EEG biomarkers across all 68 patches (*p*-values are from ANCOVA / Levene’s test of variability). Individual-subject values, mean, and SEM are shown for ASD_CA-qNL_/CON_CA-qNL_ (*blue circles*) and ASD_CA-qABN_/CON_CA-qABN_ (*red circles*). For all comparisons at the whole-brain level, *p*-values are adjusted for age, IQ, and sex as covariates.

**Figure 4.**
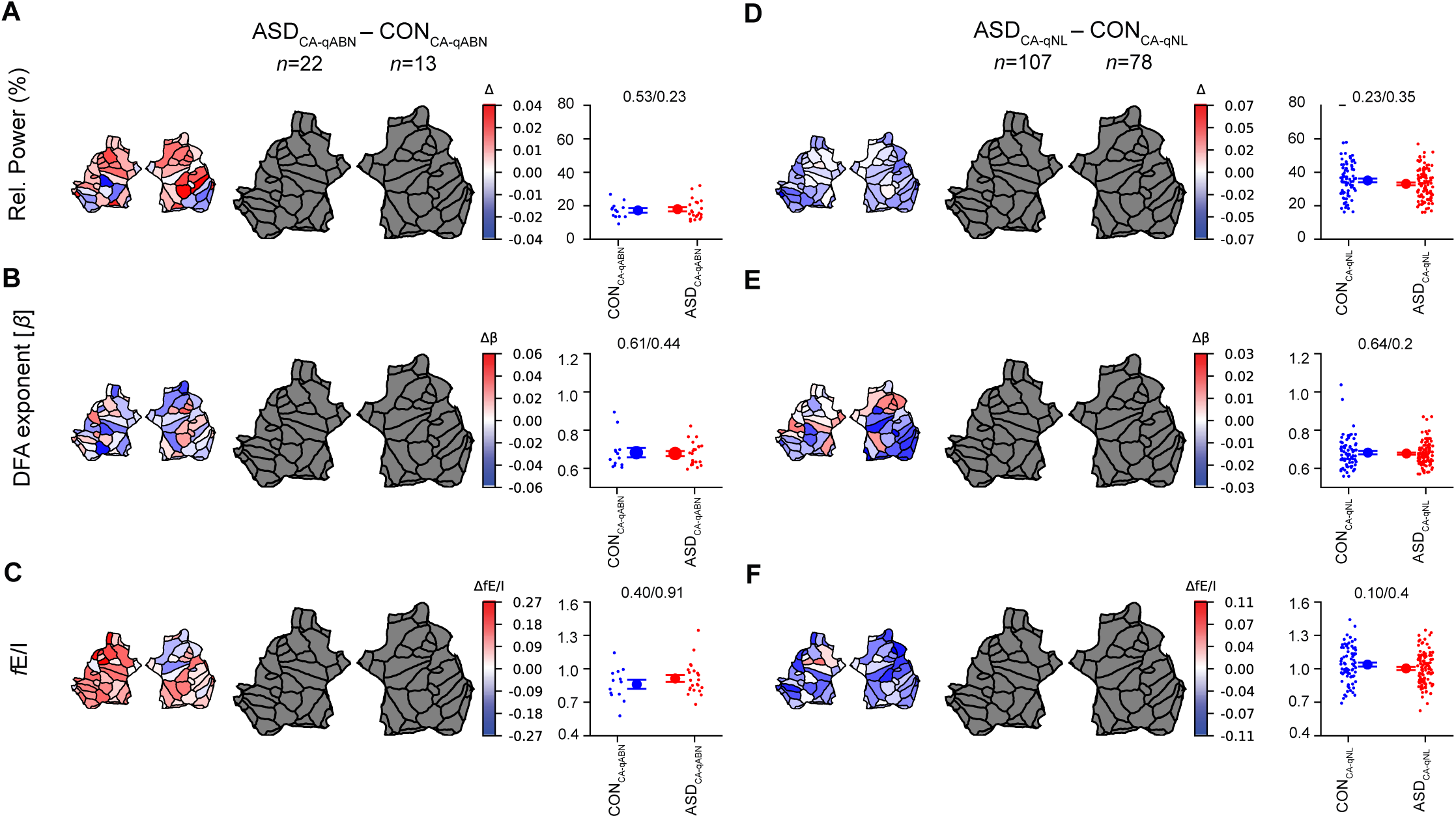
Alpha oscillations are quantitatively similar when stratifying children-adolescents with ASD and controls by qualitative EEG abnormalities. EEG measures in control children-adolescents with qualitative EEG abnormalities (CON_CA-qABN_) were not different from children-adolescents with ASD and abnormal EEG (ASD_CA-qABN_) **(A-C)**. Moreover, measures in ASD children-adolescents with normal EEG (ASD_CA-qNL_) were not different from controls with normal EEG (CON_CA-qNL_) **(D-F)**. Grand-average flatmaps are shown for the cohort difference of the indicated comparisons (labels on top), where the small flatmaps represents the mean difference, and the large plot represents significant differences (*p*-value < 0.05, Wilcoxon rank-sum test, FDR corrected) as colored patches, and insignificant differences as gray patches. Comparisons in boxplots were based on the average value of the EEG biomarkers across all 68 patches (ANCOVA / Levene’s test of variability). In the 3^rd^ column, individual-subject values, mean, and SEM are shown for CON_CA-qABN_ (*blue circles*) and ASD_CA-qABN_ (*red circles*). In the 6^th^ column, same values are shown for CON_CA-qNL_ (*blue circles*), and ASD_CA-qNL_ (*red circles*). For all comparisons at the whole brain level, *p*-values are adjusted for age, IQ, and sex as covariates.

### Children-adolescent differences between abnormal and normal EEG are robust across studies

Given the similarity of the normal vs. abnormal EEG contrasts in children-adolescents with ASD and in controls (previous section, also **Figure 3**), we grouped all children-adolescents, regardless of the presence of ASD, to further investigate the quantitative differences between those with normal vs. abnormal EEG. We source-reconstructed EEG data from a previous study on ASD children-adolescents (SPACE-BAMBI)^27^, and contrasted the biomarker values across abnormal and normal EEG. We found that the EU-AIMS and SPACE-BAMBI datasets show remarkably similar spatial contrasts patterns between normal and abnormal EEG (**Figure 5**), with both datasets showing significant patches predominantly in the right occipital, bilateral temporal and cingulate areas (**Table S7, S8**).

**Figure 5.**
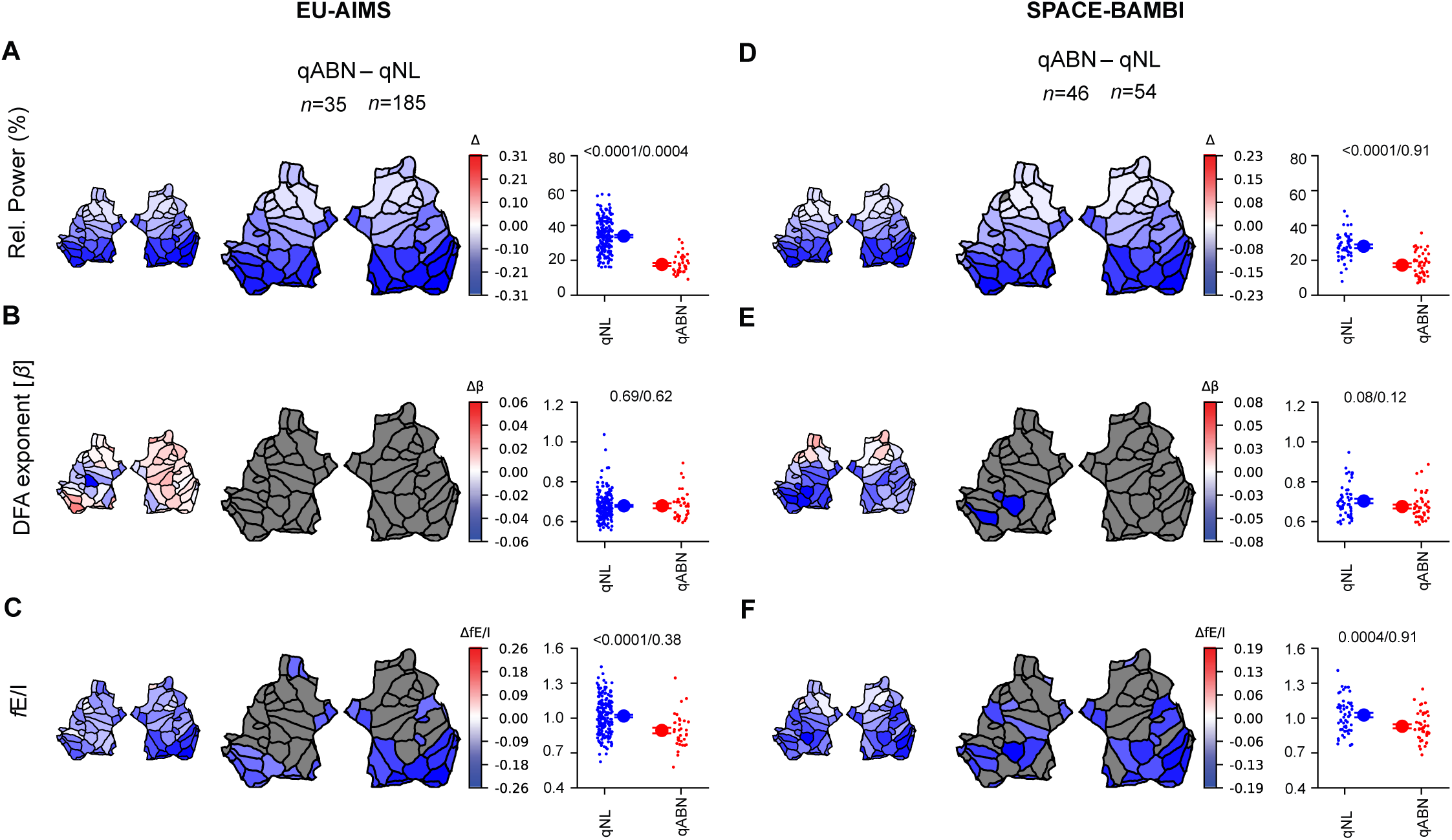
The distinction in alpha-band relative power and fE/I with respect to the presence/absence of qualitative abnormalities is robust across studies. After merging ASD and CON children-adolescent subjects that have qualitatively normal or abnormal EEG, respectively **(A-C)**, we observed very similar contrasts in alpha-band relative power and fE/I to those found in a re-analysis of the SPACE-BAMBI data previously published^27^ **(D-F)**. Grand-average flatmaps are shown for the cohort difference of the indicated comparisons (labels on top), where the small flatmaps represents the mean difference, and the large plot represents significant differences (*p*-value < 0.05, Wilcoxon rank-sum test, FDR corrected) as colored patches, and insignificant differences as gray patches. Comparisons in boxplots were based on the average value of the EEG biomarkers across all 68 patches (ANCOVA / Levene’s test of variability). For all comparisons at the whole brain level, *p*-values are adjusted for age, IQ, and sex as covariates. In the 3^rd^ and 6^th^ column, *blue* and *red circles* represent the individual-subject values, which are accompanied by the mean and SEM.

### No relationship between clinical scales and EEG biomarkers

We expected that the modulation of EEG biomarkers by the presence of EEG abnormalities would reflect in their relationship with cognitive or behavioral scales. However, we did not find a significant correlation between our set of EEG biomarkers (patch or whole-brain level) and IQ, the Autism Diagnostic Observation Scale, Repetitive Behavior Scale-Revised, or with the Vineland-II Adaptive Behavior standard score for any of the groups (ASD or controls, even for those with qualitative EEG abnormalities) (**Table S9**; Pearson correlation, FDR multiple comparisons correction).

## Discussion

The original hypothesis of E/I imbalances as a mechanism for ASD was partly based upon the observation of frequent EEG abnormalities—a hypothesis supported through diverse genetic, molecular and cellular studies^6,24,25,66–68^ but lacking support from clinical measurements. Here, we tested the implication of EEG abnormalities on E/I balance using our fE/I measure in the EU-AIMS set of EEG recordings. Before evaluating for EEG abnormalities, we found no EEG differences and equal fE/I variability between cases and controls for the children-adolescents subsamples, while the adults with ASD showed lower fE/I and relative power in the alpha band, when compared to controls. After stratifying for qualitative EEG abnormalities, we found that both children-adolescents with and without ASD and an abnormal EEG associate with low relative power and low fE/I in the alpha band, which is our main finding. While relative power effects were widespread, low fE/I in subjects with abnormal EEG was found in localized patches. Notably, EEG abnormalities in the adult population were infrequent and, therefore, we could not corroborate this EEG phenotype in the adult cohort. Overall, our results show that EEG abnormalities concur with reduced alpha-band relative power and inhibition-dominated network dynamics, in agreement with our initial study of fE/I in clinical samples^27^, and a finding we extend here by analyzing the alpha oscillations in source-reconstructed brain regions. Importantly, our findings support the notion that EEG abnormalities reflect a disturbance of E/I balance.

Our next challenge was to test whether EEG abnormalities have clinical implications or rather exist simultaneously or independently of pathology^69–71^. On average, we did not find differences in IQ or clinical scores between ASD with or without EEG abnormalities, and we did not find significant correlations between network E/I and IQ or clinical questionnaires in either the ASD cohorts or controls, even after stratification for EEG abnormalities. Indeed, the E/I ratio of cortical networks determines network function, e.g., by modulating stimulus sensitivity and long-range functional connectivity^2,3^, and specific spatial patterns of E/I imbalance are likely to result in different behavioral or cognitive deficits albeit not the ones measured with the clinical scales used in the present study, or the statistical power was insufficient. Another potential explanation is that the spectrum of ASD severity does not depend in a linear, straightforward manner, on factors that destabilize network functioning in terms of E/I balance. Furthermore, especially the lack of correlation between fE/I and clinical symptom scores in children-adolescents without EEG abnormalities may resonate with the broad clinical definition of the spectrum, in which many different mechanistic factors may contribute.

In fact, in the EU-AIMS study, EEG abnormalities were observed not only in the ASD sample but also in controls, which highlights the need for further investigation into their specificity. Indeed, EEG abnormalities are reported in relatively healthy populations, but frequently associated with common neurological conditions, e.g., migraine, visual deficits, or subclinical psychiatric disorders^55,72^ or the consequence of common medication (sleeping drugs, muscle relaxants, sedatives and antidepressants^73^. A more lenient inclusion criteria for the EU-AIMS control sample*—*which allowed for participants with different comorbidities to be enrolled and under some relevant psychotropic medication (e.g., melatonin, antidepressants and methylphenidate), may have resulted in a heterogeneous “healthy” control sample. Although we accounted for this heterogeneity to the best of our knowledge by excluding data from participants in the control group reported to use psychotropic medication or with intellectual disability, it is still plausible that some conditions or medications were not explicitly registered and thus not controlled for. In line with this, we can only speculate that the primary mechanisms underlying the EEG abnormalities found in the control group would most likely differ from those found in ASD^55–58,74–78^. In fact, the type of EEG abnormalities were different between ASD and controls, e.g., epileptiform patterns were almost exclusive to ASD while the control group mainly showed altered background activity. It remains important to explore the fE/I characteristics across different EEG abnormalities, as these are known to reflect distinct pathophysiological mechanisms involving excitatory-inhibitory systems and with different clinical repercussions. For instance, altered neuronal excitability thresholds and synchronous discharges triggered by inhibitory interneurons are associated with epileptiform interictal spikes^74,75,79^. Disrupted E-I feedback in cortical and subcortical networks can lead to rhythmic slowing^77,80^, while thalamocortical dysrhythmia has been proposed as the mechanism behind the more diffuse background slowing seen in diverse neuropsychiatric conditions^81,82^. However, this study was unable to investigate these mechanisms separately, which represents an important limitation, since we applied a broader definition of “an abnormal EEG” in order to address challenges related to the small sample size. Notably, examining E/I differences across the diverse range of EEG abnormalities could have revealed stronger associations with clinical features or differences between ASD and control groups, potentially improving the specificity of our findings. Finally, we agree that detecting qualitative EEG abnormalities through an initial qualitative EEG inspection can help establish a more rigorous level of normative criteria beforehand, and serve as a foundation for enhancing control samples^72^.

The more lenient inclusion criteria for the EU-AIMS study might also explain the lack of increased variability in fE/I in ASD, compared to controls, which was originally reported in Bruining *et al.*^27^. On the other hand, these results are consistent with other recent EEG studies, which show similar fE/I variability of ASD and controls^46,83^. This, however, does not diminish the importance of association between fE/I and the presence of EEG abnormalities, which could be of potential value for pharmacological interventions targeting restoration of E/I balance.

The association between lower fE/I and qualitative EEG abnormalities was not globalized, but consistently localized in the occipital, temporal and cingulate regions, across both the ASD and CON groups, and was replicated across the EU-AIMS and SPACE-BAMBI datasets. Differences in activity in these regions have been associated with ASD and with sensory processing in particular^84–86^. In typically developing individuals who exhibited qualitative EEG abnormalities, lower fE/I in the same areas might be related to subclinical autistic traits^87^. Additionally, the wider spread of regions with lower fE/I, in typically developing children-adolescents, when compared with autistic individuals, could reflect a more heterogeneous set of factors contributing to the qualitative abnormalities and alterations in circuitry. Future studies should aim to determine the precise clinical impact of the presence of EEG abnormalities, in ASD and typically developing individuals, beyond conventional ASD questionnaires for instance using patient report outcome measures of sensory reactivity^88^.

The lower fE/I for alpha oscillations observed in subjects with abnormal EEG is accompanied by decreased relative power in the same frequency band, which raises the question whether the fE/I reduction is a trivial effect of power shifts. However, fE/I is calculated from the correlation between windowed power and the temporal structure of amplitude fluctuations in alpha-band. Therefore, a change in fE/I could not occur only due to overall power shifts (e.g., doubling the alpha oscillation signal does not affect the correlation coefficient used to calculate fE/I), but requires a concomitant alteration in amplitude fluctuations, reflecting a different dynamical regime. Nonetheless, as has been investigated for DFA^89^, a lower alpha-band power will lead to a decrease in the signal-to-noise ratio, which eventually will interfere with measuring the actual temporal structure. When this happens, both DFA and fE/I are expected to be underestimated and this could in principle have influenced our results. We do not have a measure of signal-to-noise ratio to test this possibility; however, the absence of an effect of DFA speaks in favor of sufficient signal-to-noise ratio to adequately estimate the temporal structure and, thus, fE/I.

So, why are EEG abnormalities associated with lower fE/I? We previously studied two genetic disorders that are both strongly associated with ASD, epilepsy and intellectual disability, i.e., Tuberous Sclerosis Complex and STXBP1 syndrome^90,91^. In both disorders, patients showed reduced power and low fE/I ratios in most brain regions, when compared with healthy subjects. These findings seem counterintuitive, since epilepsy comorbidity and EEG abnormalities are traditionally considered a sign of cortical hyperexcitability. However, in an epileptogenic network, neuronal hyperexcitability can coexist with an excess of inhibition, e.g., both slowing of activity and increased localized network activity can be strong indicators of the epileptogenic zone interictally^92–95^. This interpretation would indicate a refinement of the initial E/I hypothesis in ASD and may be consistent with the notion that primary ‘local’ increases in E/I through genetic or metabolic defects, such as present in TSC or STXBP1, can trigger other compensatory inhibitory homeostatic mechanisms, e.g., synaptic scaling, lower presynaptic glutamate release and decrease postsynaptic AMPA receptor clustering or sensitivity^1,25,96^. At the network level, this may result in EEG abnormalities such as an increased slowing of activity—as it has been noted in both ASD and in the interictal period of epilepsy patients^27,49,97,98^. Furthermore, compensatory mechanisms may also account for so-called paradoxical effects that we and others observed when testing E/I-affecting drugs in ASD trials^32,99^. Together, the use of fE/I as a ‘proxy marker’ for E/I balance may help to link different neurobiological levels and reconsider oversimplistic extrapolation of cellular and local to global network-scale E/I interpretations. In clinical samples, stratification on the basis of EEG abnormalities and fE/I measurement may increase the power of studies on genetics of disease and on treatments targeting E/I balance.

## CONCLUSION

The previously published association between EEG abnormalities and lower network level functional E/I ratio in the SPACE-BAMBI cohort is replicated in ASD and controls of the EU-AIMS LEAP sample. Source modeling found strong associations between the implicated anatomical regions in the two samples. Our findings indicate a method to stratify part of the neurophysiological heterogeneity of ASD and gain understanding of the effects of homeostatic interactions in E/I balance regulation.

## Availability of data and materials

Due to privacy regulations of human subjects, we cannot provide the EEG files of the subjects included in our study. Analysis scripts to reproduce the figures and statistics, along with the underlying data, will be made available on figshare (https://figshare.com/), a website dedicated for sharing scientific data. The code for the fE/I algorithm is publicly available at https://github.com/rhardstone/fEI.

## Acknowledgements

We thank all participants and their families for participating in this study. We also gratefully acknowledge the contributions of all members of the EU-AIMS LEAP group.

## Funding

Netherlands Organization for Scientific Research (NWO) Physical Sciences Grant 612.001.123 (K.L.-H.)

Netherlands Organization for Scientific Research (NWO) Social Sciences 406-15-256 (A.-E.A.,K.L.-H.)

NWA-ORC Call (NWA.1160.18.200) (H.B., K.L.-H.).

NWO ‘BRAINMODEL’, project number 10250022110003 (2021-2027) (H.B., K.L.-H.)

BRAINinBALANCE TKI program (2022-2012377) (to H.B., K.L.-H.)

EU H2020 ‘Human Brain Project’ grant agreement no. 604102 (H.D.M.)

EU-AIMS (European Autism Interventions) and AIMS-2-TRIALS programmes which receive support from Innovative Medicines Initiative Joint Undertaking Grant No. 115300 and 777394, the resources of which are composed of financial contributions from the European Union’s FP7 and Horizon2020 programmes, and from the European Federation of Pharmaceutical Industries and Associations (EFPIA) companies’ in-kind contributions, and AUTISM SPEAKS, Autistica and SFARI; and by the Horizon2020 supported programme CANDY Grant No. 847818

## Author contributions

Conceptualization: H.B., K.L.-H., P.G., J.F.H.

Methodology: E.L. J.-M., A.-E.A., S.-S.P., K.L.-H., H.B.

Investigation: E.L.J-M, K. L-H, H.B.

Visualization: E.L. J.-M., A.-E. A.

Acquisition, analysis, or interpretation of data: All authors.

Funding acquisition: K. L.-H., H.B.

Administrative, technical, or material support: A.-E. A., S.-S.P.

Supervision: K. L-H, H.B., H.D.M.

Writing – original draft: E. L. J-M, K. L-H, H. B.

Writing – review & editing: all authors

