## Supplementary Materials for "Qualitative EEG abnormalities signal a shift towards inhibition-dominated brain networks. Results from the EU-AIMS LEAP studies"

**TABLES**

| children-adolescents | CON_CA_ (*n* = 91) | ASD_CA_ (*n* = 129) | | T (df) / Z, W | p-value |
| --- | --- | --- | --- | --- | --- |
| males/females | 57/34 | 91/38 | | – | – |
| Age  (mean ± SD) | 7–17  13.0 ± 3.0 | | 6–17  13.0 ± 3.0 | -0.1, 14228.5 | 0.96 |
| TIQ  (range, mean ± SEM) | *n* = 90  77–142  109.8 ± 1.4 | *n* = 125  54–139  98.3 ± 1.7 | | -4.4, 11536.0 | <.0001 |
| ADOS-2  (mean ± SEM) | NA | *n* = 126  10.9 ± 0.3 | | – | – |
| RBS-R  (mean ± SEM) | *n* = 78  1.1 ± 0.2 | *n* = 115  16.3 ± 1.2 | | 10.7, 15225.5 | <.0001 |
| VABS-composite  (mean ± SEM) | *n* = 36  104.4 ± 2.0 | *n* = 99  74.6 ± 1.4 | | -7.7, 5179.5 | <.0001 |
| VABS-com  (mean ± SEM) | *n* = 36  102.5 ± 2.5 | *n* = 104  78.9 ± 1.5 | | -6.5, 5970.0 | <.0001 |
| VABS-dl  (mean ± SEM | *n* = 36  101.9 ± 1.8 | *n* = 101  77.5 ± 1.7 | | -6.8, 5572.5 | <.0001 |
| VABS-soc  (mean ± SEM | *n* = 36  108.9 ± 2.1 | *n* = 102  74.2 ± 1.4 | | -13.3 (136.0) | <.0001 |
| Adults | **CON_A_ (*n* = 71)** | **ASD_A_ (*n* = 93)** | |  |  |
| males/females | 52/19 | 55/21 | |  |  |
| Age  (mean ± SD) | 18–31  22.6 ± 3.6 | 18–30  21.8 ± 3.2 | | -1.3, 7289.5 | 0.2 |
| TIQ  (range, mean ± SEM | *n* = 71  81–142  108.0 ± 1.4 | *n* = 93  59–148  99.6 ± 1.9 | | -2.8, 6824.0 | 0.005* |
| ADOS-2  (mean ± SEM) | NA | *n* = 92  10.5 ± 0.4 | | – | – |
| RBS-R  (mean ± SEM) | NA | *n* = 70  9.9 ± 1.2 | | – | – |
| VABS-composite  (mean ± SEM) | NA | *n* = 76  69.3 ± .1.7 | | – | – |
| VABS-com  (mean ± SEM) | *n* = 1 | *n* = 79  73.5 ± 2.0 | | – | – |
| VABS-dl  (mean ± SEM | *n* = 1 | *n* = 78  72.2 ± 1.8 | | – | – |
| VABS-soc  (mean ± SEM | NA | *n* = 77  70 ± 2.0 | | – | – |

**Table S1. Clinical characteristics of participants included for EEG analyses in this study**. Mean values and comparison statistics (t-test for parametric data (*t*, *df*) and Wilcoxon rank-sum test for non-parametric data (*W*, *z*). Demographics and EEG mean biomarker values per subject can be found on Table S1. CON_CA_ (control children-adolescents). CON_A_ (control adults). ASD_CA_ (children-adolescents with autism spectrum disorder). ASD_A_ (adults with autism spectrum disorder). TIQ (Total Intelligence Quotient); ADOS-2 (Autism Diagnostic Observation Scale); RBS-R (Repetitive Behavioral Scale revised); VABS (Vineland-II Adaptive Behavior standard score); VABS-composite (composite standard score); VABS-com (communication domain standard score), VABS-dl (daily living domain standard score); VABS-soc (socialization domain standard score). NA (scores not registered).

| children | con_ch_ (*n* = 35) | ASD_ch_ (*n* = 42) |  | F (DF) | p-value | p-value  levene |
| --- | --- | --- | --- | --- | --- | --- |
| $\boldsymbol{\alpha}$ relative power  (mean ± SEM) | 0.3 ± 0.017 | 0.28 ± 0.014 |  | 0.005 (69) | 0.94 | 0.52 |
| DFA  (mean ± SEM) | 0.67 ± 0.011 | 0.66 ± 0.008 |  | 1.60 (69) | 0.72 | 0.44 |
| fE/I  (mean ± SEM) | 1.03 ± 0.023 | 1.009 ± 0.021 |  | 0.13 (69) | 0.86 | 1.00 |
| adolescents | **CON_ad_ (*n* = 56)** | **ASD_ad_ (*n* = 87)** |  | **F (DF)** | **P-VALUE** | **P-VALUE**  **levene** |
| $\boldsymbol{\alpha}$ relative power  (mean ± SEM) | 0.34 ± 0.016 | 0.31 ± 0.011 |  | 0.67 (136) | 0.21 | 0.35 |
| DFA  (mean ± SEM) | 0.69 ± 0.012 | 0.69 ± 0.007 |  | 0.03 (136) | 0.41 | 0.18 |
| fE/I  (mean ± SEM) | 1.002 ± 0.024 | 0.98 ± 0.016 |  | 0.39 (136) | 0.53 | 0.15 |
| adults | **CON_A_ (*n* = 71)** | **ASD_A_ (*n* = 93)** |  | **F (DF)** | **P-VALUE** | **P-VALUE**  **levene** |
| $\boldsymbol{\alpha}$ relative power  (mean ± SEM) | 0.4 ± 0.014 | 0.35 ± 0.012 |  | 5.78 (159) | 0.02 | 0.65 |
| DFA  (mean ± SEM) | 0.69 ± 0.008 | 0.70 ± 0.008 |  | 1.28 (159) | 0.26 | 0.29 |
| fE/I  (mean ± SEM) | 1.05 ± 0.02 | 1 ± 0.019 |  | 1.18 (159) | 0.28 | 0.93 |

**Table S2. Contrast between ASD subjects and controls, across the children and adolescent age groups.** The biomarker values were averaged for each subject across all patches. Next, we compared the whole-brain averages between ASD and controls, for 3 age groups: children, adolescents and adults, using ANCOVA. *p*-values are adjusted for age, IQ, and sex as covariates.

| children-adolescents | con_cA-nl_ (*n* = 78) | con_cA-abn_ (*n* = 13) |  | T (df)/W, z | p-value |
| --- | --- | --- | --- | --- | --- |
| males/females | 48/30 | 9/4 |  | – | – |
| Age  (mean ± SD) | 7–17  13.4 ± 2.9 | 8–16  10.8 ± 2.5 |  | -2.8, 355.5 | 0.006* |
| TIQ  (range, mean ± SEM | *n* = 77  77–134  108.6 ± 1.5 | *n* = 13  96–142  116.4 ± 3.9 |  | 2.0 (88.0) | 0.05 |
| ADOS-2  (mean ± SEM) | NA | NA |  | – | – |
| RBS-R  (mean ± SEM) | *n* = 66  1 ± 0.3 | *n* = 12  1.1 ± 0.6 |  | 0.3, 494.0 | 0.78 |
| VABS-composite  (mean ± SEM) | *n* = 30  102.5 ± 2.2 | *n* = 6  114.3 ± 3.2 |  | 2.3 (34) | 0.027* |
| VABS-com  (mean ± SEM) | *n* = 30  100 ± 2.5 | *n* = 6  115 ± 6.9 |  | 2.4 (34) | 0.024* |
| VABS-dl  (mean ± SEM | *n* = 30  100.7 ± 2 | *n* = 6  107.8 ± 3.4 |  | 1.5 (34) | 0.14 |
| VABS-soc  (mean ± SEM | *n* = 30  107.2 ± 2.3 | *n* = 6  117.5 ± 2.6 |  | 1.9 (34) | 0.06 |
| children-adolescents | **ASD_CA-qNL_ (*n* = 107)** | **ASD_CA-qABN_ (*n* = 22)** |  |  |  |
| males/females | 83/24 | 8/14 |  | – | – |
| Age  (mean ± SD) | 7–17  13.3 ± 2.9 | 6–17  11.5 ± 3.0 |  | -2.3, 1055.5 | 0.019* |
| TIQ  (range, mean ± SEM | *n* = 105  54–139  98.9 ± 1.9 | *n* = 20  55–124  95.5 ± 4.7 |  | -0.6, 1168.0 | 0.54 |
| ADOS-2  (mean ± SEM) | *n* = 105  10.8 ± 0.4 | *n* = 21  11.3 ± 0.8 |  | 0.7, 1436.5 | 0.5 |
| RBS-R  (mean ± SEM) | *n* = 94  16.6 ± 1.4 | *n* = 21  15.1 ± 1.7 |  | 0.4, 1266.5 | 0.73 |
| VABS-composite  (mean ± SEM) | *n* = 83  75.1 ± 1.5 | *n* = 16  72.1 ± 2.9 |  | -0.9, 704.0 | 0.36 |
| VABS-com  (mean ± SEM) | *n* = 86  78.8 ± 1.7 | *n* = 18  79.2 ± 3.9 |  | -0.1, 936.0 | 0.94 |
| VABS-dl  (mean ± SEM | *n* = 85  78.1 ± 1.9 | *n* = 16  74.2 ± 3.7 |  | -1.1, 702.0 | 0.29 |
| VABS-soc  (mean ± SEM | *n* = 84  75.1 ± 1.5 | *n* = 18  70.1 ± 2.8 |  | -1.2, 787.5 | 0.22 |

**Table S3. Clinical differences between children-adolescents with and without qualitative EEG abnormalities in the control and the ASD groups.** Mean values and comparison statistics (t-test for parametric data (*t*, *df*) and Wilcoxon rank-sum test for non-parametric data (*W*, *z*). CON_CA-_q_NL_ (control children-adolescents with normal EEG). CON_CA-qABN_ (control children-adolescents with EEG abnormalities). ASD_CA-NL_ (children-adolescents with ASD with normal EEG). ASD_CA-qABN_ (children-adolescents with ASD and EEG abnormalities). TIQ (Total Intelligence Quotient); ADOS-2 (Autism Diagnostic Observation Scale); RBS-R (Repetitive Behavioral Scale revised); VABS (Vineland-II Adaptive Behavior standard score); VABS-composite (composite standard score); VABS-com (communication domain standard score), VABS-dl (daily living domain standard score); VABS-soc (socialization domain standard score). *Significant *p-*values (< 0.05).

| **Region** | **Significant patches** | **Total patches** | **% significant patches** |
| --- | --- | --- | --- |
| *Occipital_rh* | *4* | *4* | *100* |
| *Cingulate_rh* | *1* | *4* | *25* |
| *Temporal_rh* | *2* | *9* | *22* |
| *Frontal_rh* | *1* | *11* | *9* |
| Frontal_lh | 0 | 11 | 0 |
| Temporal_lh | 0 | 9 | 0 |
| Parietal_lh | 0 | 5 | 0 |
| Parietal_rh | 0 | 5 | 0 |
| Occipital_lh | 0 | 4 | 0 |
| Cingulate_lh | 0 | 4 | 0 |
| Insular_lh | 0 | 1 | 0 |
| Insular_rh | 0 | 1 | 0 |

**Table S4. Percentage of significant patches in each region, for the contrast ASD_CA-qABN_** **- ASD_CA-qNL_.** Regions were sorted by the % of significant patches. Rh and lh correspond to the right and left hemispheres.

| **Region** | **Significant patches** | **Total patches** | **% significant patches** |
| --- | --- | --- | --- |
| *Occipital_lh* | *4* | *4* | *100* |
| *Occipital_rh* | *4* | *4* | *100* |
| *Insular_rh* | *1* | *1* | *100* |
| *Parietal_rh* | *3* | *5* | *60* |
| *Temporal_lh* | *5* | *9* | *56* |
| *Temporal_rh* | *5* | *9* | *56* |
| *Cingulate_lh* | *2* | *4* | *50* |
| *Cingulate_rh* | *1* | *4* | *25* |
| *Frontal_lh* | *2* | *11* | *18* |
| Frontal_rh | 0 | 11 | 0 |
| Parietal_lh | 0 | 5 | 0 |
| Insular_lh | 0 | 1 | 0 |

**Table S5. Percentage of significant patches in each region, for the contrast CON_CA-qABN_** **- CON_CA-qNL_.** Regions were sorted by the % of significant patches. Rh and lh correspond to the right and left hemispheres.

| children-adolescents | con_CA-qNL_ (*n* = 78) | ASD_CA-qNL_ (*n* = 107) |  | F (DF) | p-value | p-value  levene |
| --- | --- | --- | --- | --- | --- | --- |
| $\boldsymbol{\alpha}$ relative power  (mean ± SEM) | 0.35 ± 0.011 | 0.33 ± 0.009 |  | 1.43 (177) | 0.23 | 0.35 |
| DFA  (mean ± SEM) | 0.68 ± 0.008 | 0.68 ± 0.006 |  | 0.23 (177) | 0.64 | 0.2 |
| fE/I  (mean ± SEM) | 1.04 ± 0.018 | 1.003 ± 0.014 |  | 2.70 (177) | 0.10 | 0.4 |
| children-adolescents | **CON_CA-qABN_ (*n* = 13)** | **ASD_CA-qABN_ (*n* = 22)** |  | **F (DF)** | **P-VALUE** | **P-VALUE**  **levene** |
| $\boldsymbol{\alpha}$ relative power  (mean ± SEM) | 0.17 ± 0.013 | 0.18 ± 0.013 |  | 0.40 (28) | 0.53 | 0.23 |
| DFA  (mean ± SEM) | 0.68 ± 0.025 | 0.68 ± 0.013 |  | 0.26 (28) | 0.61 | 0.44 |
| fE/I  (mean ± SEM) | 0.86 ± 0.004 | 0.91 ± 0.03 |  | 0.73 (28) | 0.40 | 0.91 |

**Table S6. Contrast between subjects ASD and controls, across the groups with and without qualitative EEG abnormalities.** The biomarker values were averaged for each subject across all patches. Next, we compared the whole-brain averages of EEG biomarkers, between ASD and controls, for children-adolescents with and without qualitative EEG abnormalities, using ANCOVA. *p*-values are adjusted for age, IQ, and sex as covariates.

| **Region** | **Significant patches** | **Total patches** | **% significant patches** |
| --- | --- | --- | --- |
| *Occipital_lh* | *4* | *4* | *100* |
| *Occipital_rh* | *4* | *4* | *100* |
| *Insular_rh* | *1* | *1* | *100* |
| *Temporal_rh* | *7* | *9* | *78* |
| *Cingulate_lh* | *3* | *4* | *75* |
| *Cingulate_rh* | *2* | *4* | *50* |
| *Temporal_lh* | *4* | *9* | *44* |
| *Parietal_rh* | *2* | *5* | *40* |
| *Frontal_lh* | *2* | *11* | *18* |
| *Frontal_rh* | *1* | *11* | *9* |
| Parietal_lh | 0 | 5 | 0 |
| Insular_lh | 0 | 1 | 0 |

**Table S7. Percentage of significant patches in each region, for the contrast qABN-qNL in the EU-AIMS dataset.** Regions were sorted by the % of significant patches. Rh and lh correspond to the right and left hemispheres.

| **Region** | **Significant patches** | **Total patches** | **% significant patches** |
| --- | --- | --- | --- |
| *Occipital_rh* | *4* | *4* | *100* |
| *Insular_lh* | *1* | *1* | *100* |
| *Insular_rh* | *1* | *1* | *100* |
| *Cingulate_rh* | *3* | *4* | *75* |
| *Parietal_lh* | *3* | *5* | *60* |
| *Temporal_rh* | *5* | *9* | *56* |
| *Cingulate_lh* | *2* | *4* | *50* |
| *Temporal_lh* | *4* | *9* | *44* |
| *Parietal_rh* | *2* | *5* | *40* |
| *Occipital_lh* | *1* | *4* | *25* |
| *Frontal_lh* | *1* | *11* | *9* |
| *Frontal_rh* | *1* | *11* | *9* |

**Table S8. Percentage of significant patches in each region, for the contrast qABN-qNL in the SPACE-BAMBI dataset.** Regions were sorted by the % of significant patches. Rh and lh correspond to the right and left hemispheres.

| children-adolescents | con_cA-qnl_ (*n* = 78) | con_cA-qabn_ (*n* = 13) | ASD_cA-qNL_ (*n* = 107) | ASD_cA-qABN_ (*n* = 22) |
| --- | --- | --- | --- | --- |
| IQ | | | | |
| $\boldsymbol{\alpha}$ relative power | *r*= -0.07, *p*=1 | *r*= -0.61, *p*=0.83 | *r*= 0.11, *p*=0.83 | *r*= 0.26, *p*=0.83 |
| DFA | *r*= -0.17, *p*=0.83 | *r*= -0.35, *p*=0.83 | *r*= -0.09, *p*=1 | *r*= 0.01, *p*=1 |
| fEI | *r*= -0.06, *p*=1 | *r*= 0.02, *p*=1 | *r*= 0.01, *p*=1 | *r*= 0.40, *p*=0.83 |
| ADOS-2 | | | | |
| $\boldsymbol{\alpha}$ relative power | – | – | *r*= 0.03, *p*=1 | *r*= -0.01, *p*=1 |
| DFA | – | – | *r*= -0.04, *p*=1 | *r*= 0.18, *p*=1 |
| fEI | – | – | *r*= 0.16, *p*=0.83 | *r*= 0.05, *p*=1 |
| RBS-R | | | | |
| $\boldsymbol{\alpha}$ relative power | *r*= -0.26, *p*=0.83 | *r*= -0.1, *p*=1 | *r*= -0.13, *p*=0.83 | *r*= 0.16, *p*=1 |
| DFA | *r*= -0.20, *p*=0.83 | *r*= -0.33, *p*=0.83 | *r*= 0.11, *p*=0.83 | *r*= -0.17, *p*=1 |
| fEI | *r*= -0.07, *p*=1 | *r*= -0.14, *p*=1 | *r*= 0.01, *p*=1 | *r*= 0.10, *p*=1 |
| VABS-composite | | | | |
| $\boldsymbol{\alpha}$ relative power | *r*= -0.26, *p*=0.83 | *r*= -0.26, *p*=1 | *r*= 0.02, *p*=1 | *r*= 0.13, *p*=1 |
| DFA | *r*= -0.32, *p*=0.83 | *r*= 0.54, *p*=0.83 | *r*= -0.03, *p*=1 | *r*= -0.28, *p*=0.83 |
| fEI | *r*= -0.27, *p*=0.83 | *r*= -0.09, *p*=1 | *r*= 0.07, *p*=1 | *r*= 0.22, *p*=1 |

**Table S9. Correlations of whole-brain averaged EEG biomarkers and clinical scales.** The biomarker values were averaged for each subject across all patches, and correlated with the scores from ADOS-2 (Autism Diagnostic Observation Scale); RBS-R (Repetitive Behavioral Scale revised); VABS (Vineland-II Adaptive Behavior standard score); VABS-composite (composite standard score), using Spearman correlation, differentiating ASD from controls, and by presence of qualitative abnormalities in the EEG. P-values were corrected using FDR with *q* = 0.05, across all correlations.

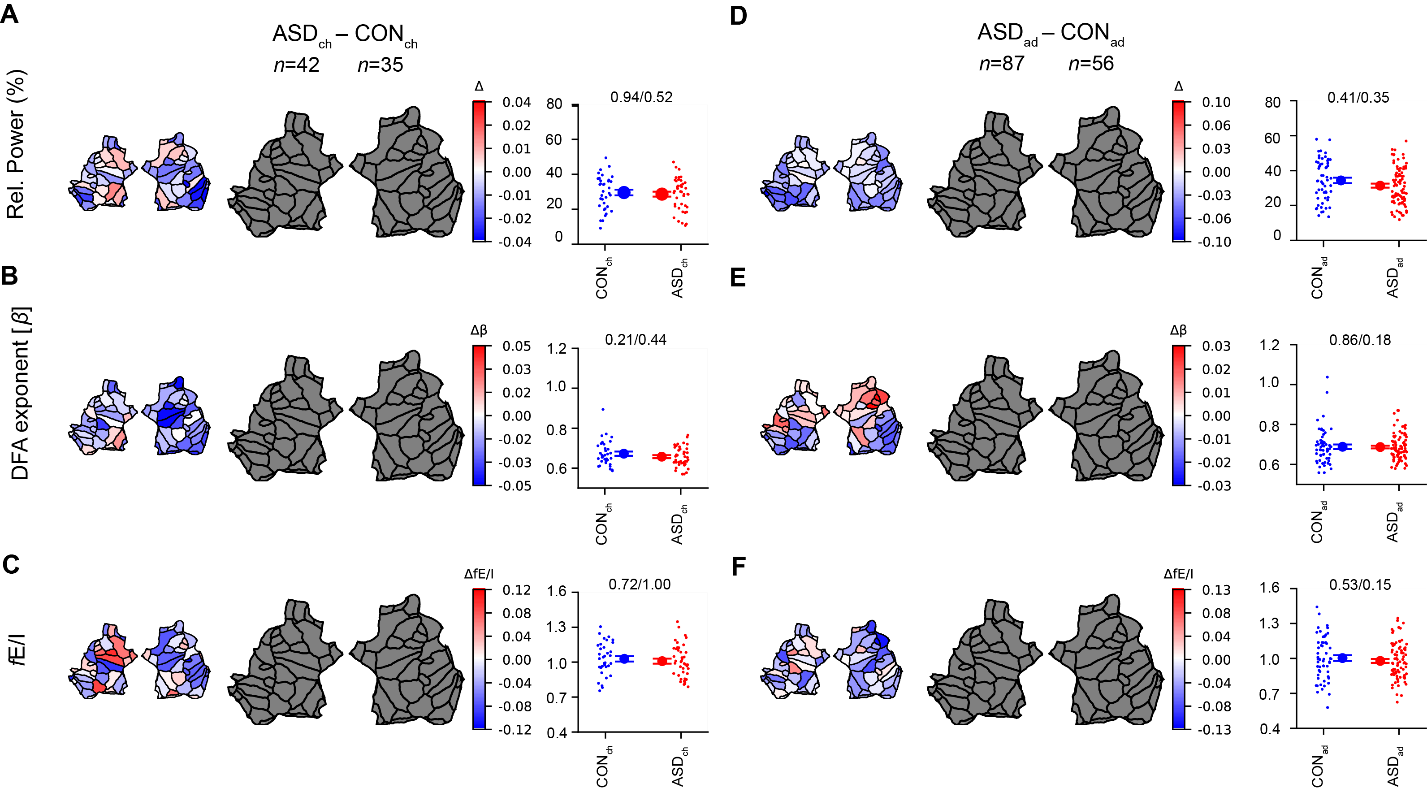

**Figure S1. Neither children nor adolescents show a significant contrast between ASD and CON.** After separating the subjects into a group of children (6–11 years) (**A-C**), and a group of adolescents (12–17 years), we did not observe any significant differences between ASD subjects and controls. Grand-average flatmaps are shown for the cohort difference of the indicated comparisons (labels on top), where the small flatmaps represents the mean difference, and the large plot represents significant differences (*p*-value < 0.05, Wilcoxon rank-sum test, FDR corrected) as colored patches, and insignificant differences as gray patches. Whole-brain difference was computed as the average over EEG biomarkers across all 68 patches (ANCOVA / Levene’s test of variability). For all comparisons at the whole-brain level, *p*-values are adjusted for age, IQ, and sex as covariates.

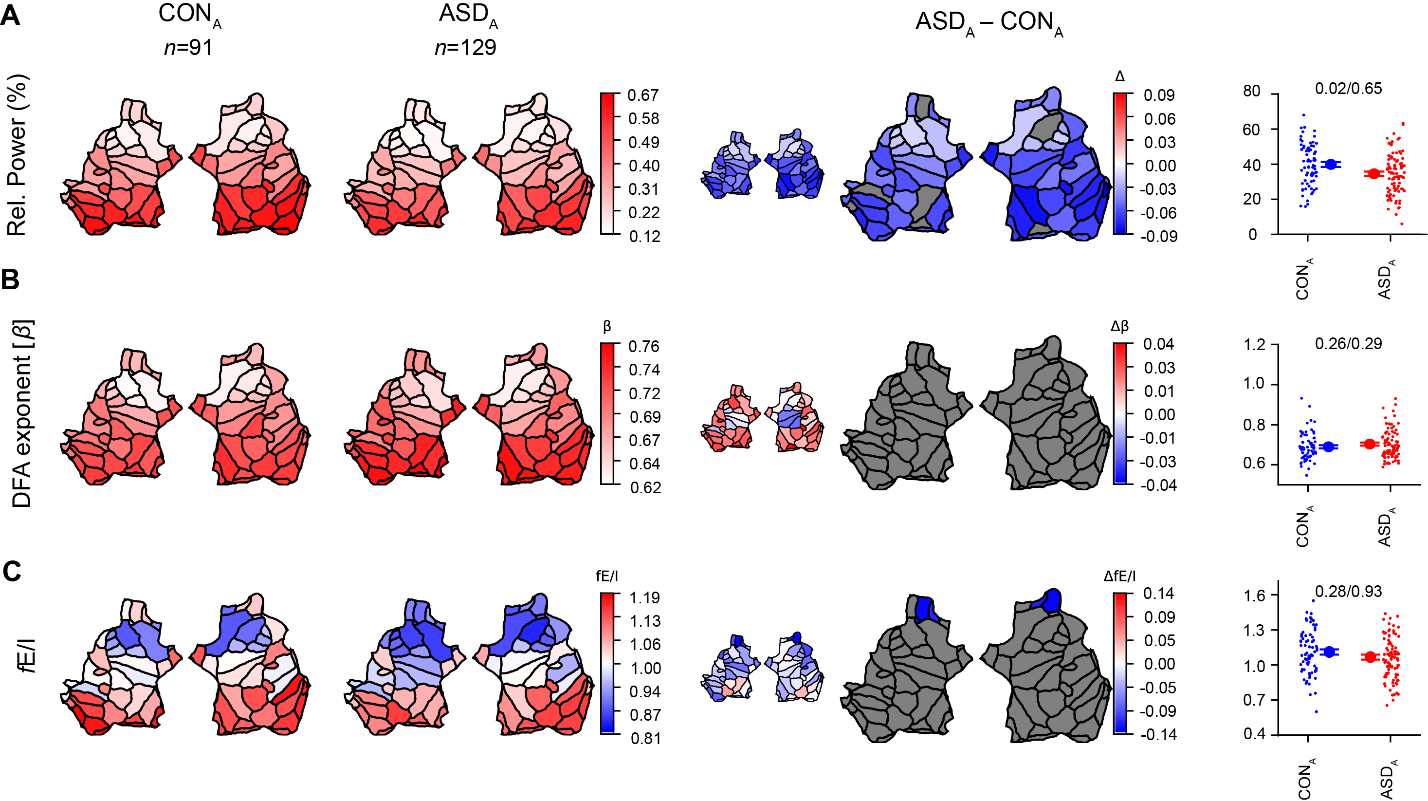

**Figure S2. Adults with ASD show significantly reduced alpha-band relative power and fE/I when compared with CON.** A majority of the patches showed a significant reduction in relative power (FDR corrected in adults with ASD when compared with adult controls (A). At a whole-brain level, however, the differences are not significant. (C) fEI shows reductions isolated in 4 frontal patches (FDR corrected). Whole-brain differences are not significant. Grand-average flatmaps are shown for the cohort difference of the indicated comparisons (labels on top), where the small flatmaps represents the mean difference, and the large plot represents significant differences (*p*-value < 0.05, Wilcoxon rank-sum test, FDR corrected) as colored patches, and insignificant differences as gray patches. Whole-brain difference was computed as the average over EEG biomarkers across all 68 patches (ANCOVA / Levene’s test of variability). For all comparisons at the whole-brain level, *p*-values are adjusted for age, IQ, and sex as covariates.
